# Ketogenesis Impact on Liver Metabolism Revealed by Proteomics of Lysine β-hydroxybutyrylation

**DOI:** 10.1101/2021.01.21.427645

**Authors:** Kevin B. Koronowski, Carolina M. Greco, He Huang, Jin-Kwang Kim, Jennifer L. Fribourgh, Priya Crosby, Carrie L. Partch, Feng Qiao, Yingming Zhao, Paolo Sassone-Corsi

## Abstract

Ketone bodies are evolutionarily conserved metabolites that function as energy substrates, signaling molecules and epigenetic regulators. β-hydroxybutyrate (β-OHB) is utilized in lysine β-hydroxybutyrylation (Kbhb) of histones, which associates with starvation-responsive genes, effectively coupling ketogenic metabolism with gene expression. The emerging diversity of the lysine acylation landscape prompted us to investigate the full proteomic impact of Kbhb. Global protein Kbhb is induced in a tissue-specific manner by a variety of interventions that evoke β-OHB. Mass spectrometry analysis of the β-hydroxybutyrylome in mouse liver revealed 891 sites of Kbhb within 267 proteins enriched for fatty acid, amino acid, detoxification and 1-carbon metabolic pathways. Kbhb of S-adenosyl-L-homocysteine hydrolase (AHCY), a rate-limiting enzyme of the methionine cycle, results in inhibition of enzymatic activity. Our results illuminate the role of Kbhb on hepatic metabolism under ketogenic conditions and demonstrate the functional consequence of this modification on a central metabolic enzyme.

## INTRODUCTION

Ketogenesis is an evolutionarily conserved mechanism that provides metabolic intermediates and alternative fuel sources (Puchalska and Crawford, 2017),(Cahill, 2006). In the liver, fatty acid oxidation-derived acetyl-CoA produces β-hydroxybutyrate (β-OHB), acetoacetate and acetone, three primary ketone bodies that are circulated among extrahepatic tissues and metabolized. This process is leveraged during periods of high fatty acid oxidation and diminished carbohydrate availability, as is evident during the neonatal period, fasting, starvation, prolonged exercise, and adherence to a ketogenic diet, where it can become a major contributor to whole organismal metabolism (Koeslag et al., 1980; McGarry and Foster, 1980) (Balasse and Fery, 1989). Physiological ketone body concentration in the blood ranges from 100 µM to 250 µM, though levels increase to ∼1 mM with a 24 hr fast and can reach ∼20 mM in pathological states like unchecked diabetes ^(Balasse and Fery, 1989; Cahill, 2006)^. The most abundant ketone body, β-OHB, is particularly intriguing given its diverse bioactive properties. While the predominant fate of β-OHB is terminal oxidation as an energy substrate, a wealth of evidence demonstrates its implication in cellular signaling, epigenetic control and posttranslational modification (PTM) of histone lysines (Puchalska and Crawford, 2017) (Xie et al., 2016) (Ruan and Crawford, 2018).

Lysine acetylation is directly linked to cellular metabolism (Peleg et al., 2016) (Katada et al., 2012). Levels of acetyl-CoA, along with acetyltransferase and deacetylase enzymes, dynamically regulate protein acetylation in response to changes in metabolic flux through corresponding pathways, for example the TCA cycle and fatty acid oxidation (Choudhary et al., 2009) (Verdin and Ott, 2015). Other Acyl-CoA metabolites, many of which are key intermediates in cellular metabolism, support a diversity of acyl-lysine modifications, including succinylation, malonylation, glutarylation, crotonylation and others (Wagner and Hirschey, 2014) (Ringel et al., 2018) (Sabari et al., 2015) (Rardin et al., 2013a). As alluded to, β-OHB serves as a substrate for histone lysine β-hydroxybutyrylation (Kbhb) via its activated thioester form β-hydroxybutyryl-CoA (Xie et al., 2016). In starved mice, increased β-OHB levels result in Kbhb on many histone lysine residues in the liver. Specifically, H3K9bhb is associated with a set of starvation-responsive genes that are distinct from genes associated with acetylation at the same H3K9 residue (Xie et al., 2016). Thus, Kbhb appears to facilitate the adaptive response to starvation by coupling metabolism to gene expression.

Lysine acylation occurs on a large variety of non-histone proteins, adding another layer of regulation to diverse cellular and metabolic pathways(Carrico et al., 2018; Lin et al., 2012). Characterization of the proteome-wide lysine β-hydroxybutyrylome is an essential step to define Kbhb cellular targets and enable investigation of the impact of ketogenesis at the cellular level. Here, we show the induction of global protein Kbhb in the liver and kidney under various conditions of elevated β-OHB. We identified 891 sites of Kbhb within 267 proteins in starved liver. These belong to macronutrient, detoxification and 1-carbon metabolic pathways, among others. Furthermore, we demonstrate that Kbhb directly inhibits the activity of the rate-limiting methionine cycle enzyme S-adenosyl-L-homocysteine hydrolase (AHCY). Our results reveal the full landscape of Kbhb and demonstrate a regulatory mechanism of Kbhb on a key metabolic enzyme under ketogenesis, implicating Kbhb in physiological and pathological processes.

## RESULTS

### Global protein β-hydroxybutyrylation *in vivo* and *in vitro*

To assess the extent of Kbhb on cellular metabolism, a pan-β-hydroxybutyryllysine (Kbhb) antibody was used to probe a panel of mouse tissues under various physiological conditions. As reported previously, starvation (48 hr fasting) elevated blood β-OHB and increased histone Kbhb in the liver (Figure 1A and S1A). Remarkably, whole-cell lysates showed a similarly significant induction of Kbhb signal for proteins of various molecular weights (Figure 1B) (Xie et al., 2016). Probing subcellular fractions revealed that Kbhb was induced broadly on cytosolic, mitochondrial and nuclear proteins (Figure 1C). As β-OHB is released from the liver and metabolized throughout the body, we probed other metabolic tissues. Starvation markedly induced Kbhb in the kidney but not pancreas, skeletal muscle, colon, heart or cerebral cortex (Figure 1D), suggesting that tissue-specific metabolism of β-OHB influences Kbhb. To assess the extent of protein Kbhb *in vitro*, cultured cells were treated with sodium-β-OHB for 24 hr. Mouse HEP1C, MEF and human HEK293T cells displayed concentration-dependent increases in protein Kbhb in response to treatment (Figure 1E). Together, these data demonstrate global protein Kbhb.

**Figure 1.**
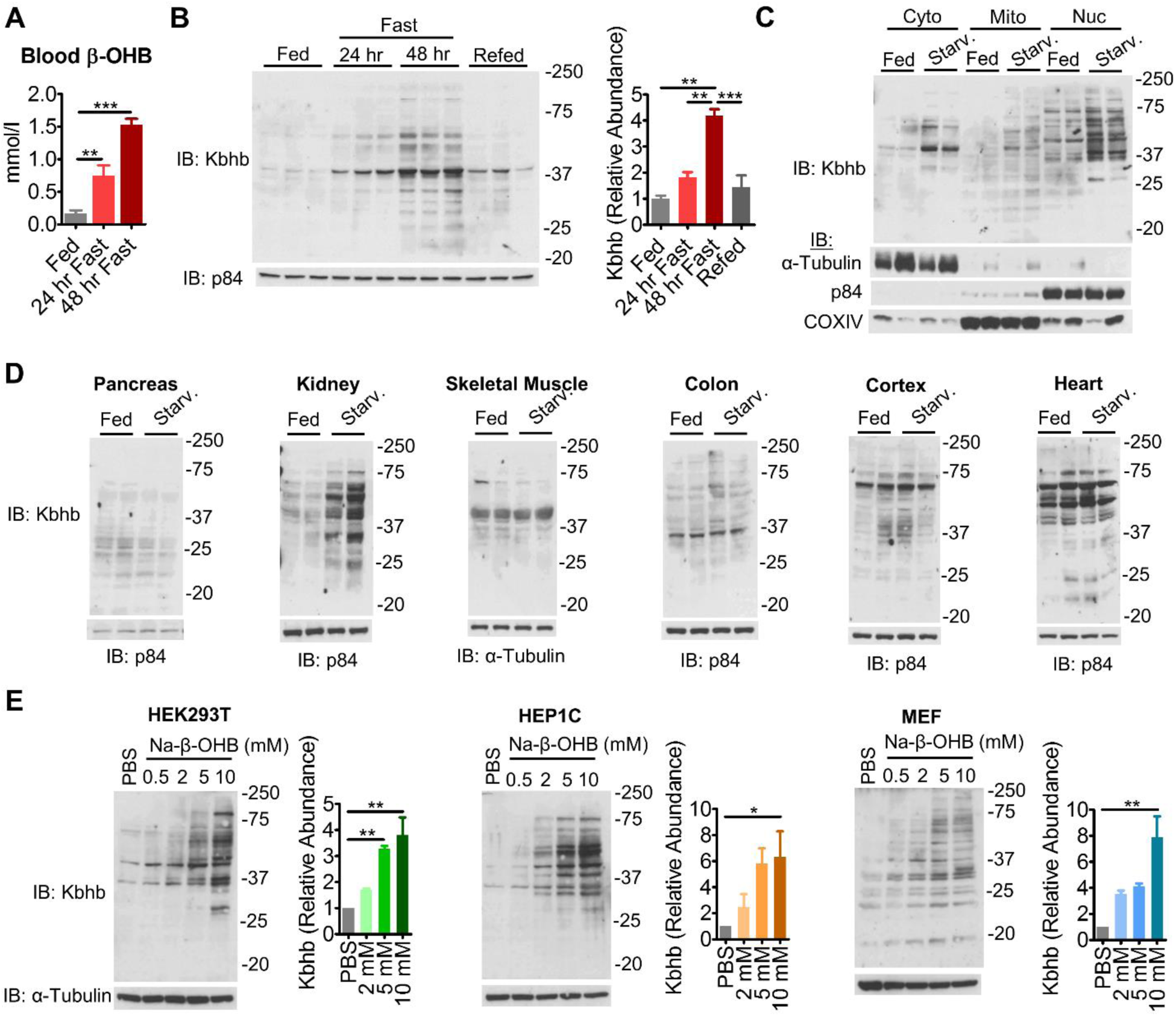
β-hydroxybutyrate is linked to global protein Kbhb *in vivo* and *in vitro*. (A) Blood β-hydroxybutyrate (β-OHB) levels in fed vs starved (48 hr fast) mice. n=3-6, unpaired Student’s t-test, ***=p<0.001. (B) Western blot for pan-β-hydroxybutyryl-lysine (Kbhb) from liver whole cell lysates. n=3 replicates are quantified to the right, unpaired Student’s t-test, **=p<0.01 **c**, Western blot from livers fractionated into cytosolic (Cyto), mitochondrial (Mito) and nuclear (Nuc) compartments. Fraction enrichment is demonstrated by compartment-specific loading controls. (D) Whole cell lysates were prepared from metabolic tissues to probe Kbhb by western blot. (E) Representative western blots from cultured cell lines treated with dose response amounts of sodium-β-hydroxybutyrate (Na-β-OHB) for 24 hr. n=3 replicates are quantified to the right, One-way ANOVA, *=p<0.05, **=p<0.01.

### Ketogenic conditions and β-OHB induce protein β-hydroxybutyrylation

Assuming that Kbhb is a general consequence of ketogenesis, then various conditions that evoke elevation of β-OHB production should also evoke Kbhb. We placed mice on a ketogenic diet (KD) for 4 weeks, which elicited ketogenesis and increased β-OHB (Figure 2A) (Newman et al., 2017) (Sato et al., 2017; Tognini et al., 2017). KD stimulated global protein Kbhb in the liver and kidney, but not other metabolic tissues, as observed under starvation (Figure 2A-B). Since KD induces circadian oscillations of β-OHB(Smith and Robinson, 2016), we collected samples along the circadian cycle and observed no apparent changes in Kbhb levels in Western analyses for the proteins detected under these experimental conditions (Figure S2A).

**Figure 2.**
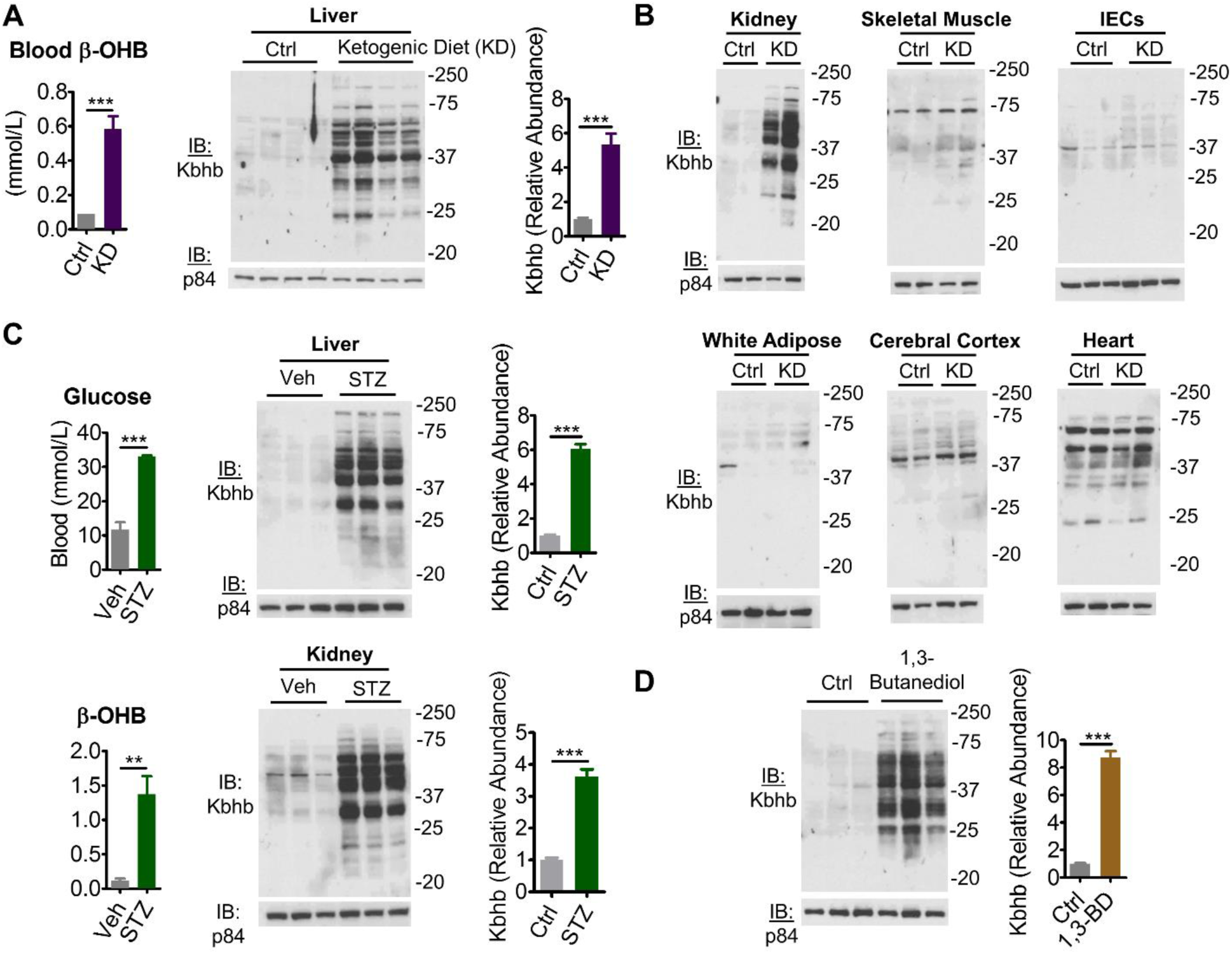
Various ketogenic conditions present with global protein β-hydroxybutyrylation. (A) left - blood β-hydroxybutyrate (β-OHB) levels in mice fed control (Ctrl) or ketogenic diet (KD) for 4 wk. n=6, unpaired Student’s t-test, ***=p<0.001; right - Western blot for pan-β-hydroxybutyryl-lysine (Kbhb) from liver whole cell lysates. n=4 replicates are quantified to the right, unpaired Student’s t-test, ***=p<0.001. (B) Whole cell lysates from Ctrl or KD conditions of the indicated tissue. IECs – intestinal epithelial cells. (C) Type I diabetes mellitus was induced by IP injection of vehicle (Veh) or 200 mg/kg streptozotocin (STZ). Blood measurements taken (left) and tissue whole cell lysates (right) prepared 4 d post injection. n=3 replicates for liver are quantified. Unpaired Student’s t-test, **=p<0.01, ***=p<0.01. (D) Liver whole cell lysates from mice fed control (ctrl) or 10% w/w 1,3-butanediol (1,3-BD) diet were probed by western blot. n=3 replicates are quantified, unpaired Student’s t-test, ***=p<0.001.

Uncontrolled type I diabetes mellitus (TIDM) is a disease state that induces ketogenesis and presents with ketoacidosis(Ruan and Crawford, 2018). Mice were administered streptozotocin (STZ), which induces TIDM by β cell destruction (Furman, 2015), evidenced by hyperglycemia coinciding with an elevation of blood β-OHB (Figure 2C). Liver and kidney from STZ-treated mice displayed a marked increase of global protein Kbhb compared to the control treatment (Figure 2C).

To determine if β-OHB alone is sufficient for Kbhb, we turned to 1,3-butanediol. This ketone precursor is converted to β-OHB by the hepatic alcohol dehydrogenase system and can be administered orally to increase β-OHB (Hashim and VanItallie, 2014; Veech, 2014). Livers from mice fed for 3 weeks a diet 10% (w/w) 1,3-butanediol displayed a marked induction of protein Kbhb (Figure 2D). Together, these data demonstrate that dietary interventions, disease states and β-OHB precursors evoke global protein Kbhb.

### Characterizing the lysine β-hydroxybutyrylome in mouse liver

We sought to identify the specific proteins and sites of Kbhb *in vivo*. To do so, we enriched Kbhb-modified peptides from starved mouse liver for analysis by mass spectrometry (MS) (Figure 3A). Proteins from whole-cell lysates were trypsin digested and the resulting peptides were fractionated by HPLC to increase coverage depth. β-hydroxybutyrylated peptides were then isolated by immunoaffinity enrichment using a polyclonal pan-Kbhb antibody (Xie et al., 2016). Samples were analyzed by liquid chromatography (LC)-MS/MS on an Orbitrap Velos MS and data were mapped to the mouse proteome. Using this method, we identified 891 sites of Kbhb across 267 proteins (Figure 3B; Table S1). Examining their subcellular distribution revealed a widespread cellular impact, as all major compartments were represented (Figure 3B).

**Figure 3.**
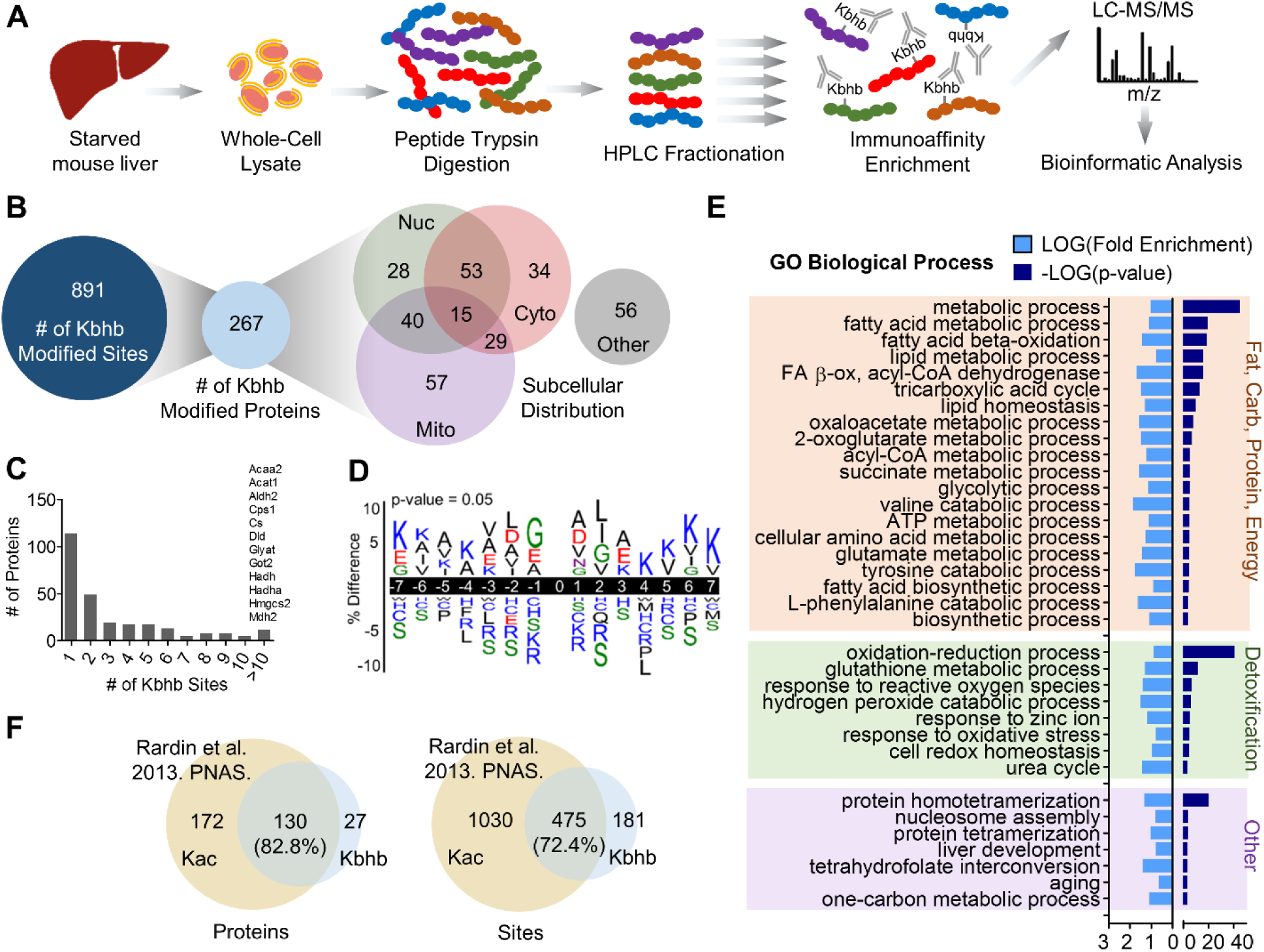
Characterizing the lysine β-hydroxybutyrylome in mouse liver. (A) Schematic of β-hydroxybutyryl-lysine (Kbhb) modified peptide identification. LC-liquid chromatography; MS-mass spectrometry. (B) The number of Kbhb sites, proteins and their subcellular distribution as determined by the COMPARTMENTS database. (C) Histogram of the number of Kbhb sites per protein. Proteins with more than 10 sites of Kbhb are listed above the corresponding bar. (D) Sequence logo of Kbhb sites determined by iceLogo. Amino acids are colored by side chain properties: blue – positively charged; red – negatively charged; green – polar uncharged; black – hydrophobic. 0=modification site. (E) Results from DAVID Gene Ongology (GO) analysis. Significantly enriched pathways (p<0.001) were categorized by cellular function for visualization. (F) Overlap of Kbhb proteins and sites with Kac proteins and sites from Rardin et al, 2014. PNAS (24 hr starved mouse liver mitochondrial fractions). Only mitochondrial proteins (determined by MitoMiner) were compared.

The number of sites per protein varied, with the majority (114, 42.7%) having one site (Figure 3C). However, several key metabolic enzymes were heavily β-hydroxybutyrylated at more than 10 sites. For example, CPS1, a rate-limiting enzyme of the urea cycle, and HMGCS2, the rate-limiting enzyme of ketogenesis, contained 35 and 15 sites of Kbhb, respectively. To determine a possible consensus motif for Kbhb, we compared the amino acid sequences surrounding the modification sites using iceLogo (Colaert et al., 2009). Positions close to the modified site (−2 to +2) were modestly enriched with glycine (small and flexible) and hydrophobic side chain-containing amino acids (leucine, alanine and isoleucine) whereas the positively charged amino acids lysine and arginine, as well as serine (polar uncharged side chain) were excluded (Figure 3D).

To understand the potential regulation by Kbhb on liver function, we performed Gene Ontology (GO) analysis using DAVID (Huang da et al., 2009). Numerous metabolic networks central to the liver were enriched with Kbhb-modified proteins, nevertheless energy and detoxification pathways were the most apparent (Figure 3E). Nearly all macronutrient pathways were top hits, encompassing fatty acid (lipid and acyl-CoA metabolic process, β-oxidation), TCA cycle, glycolytic/gluconeogenic, amino acid (several), ketone body (3 of 4 enzymes) and ATP metabolic processes. During prolonged fasting, several of these pathways are metabolizing ketogenic substrates such as fatty acid-derived acetyl-CoA and ketogenic amino acids into ketone bodies (Puchalska and Crawford, 2017), thus Kbhb appears to feedback onto the biosynthetic processes that induce it. Cellular detoxification pathways were also enriched, specifically glutathione metabolism, response to reactive oxygen species and oxidative stress, hydrogen peroxide catabolism and redox homeostasis. Other notable enrichment included nucleosome assembly, evidenced by identification of Kbhb on various histone species, as well as cofactor pathways such as tetrahydrofolate interconversion and one-carbon metabolic process. Notably, the one-carbon cycle and methionine cycle constitute a key metabolic node in the liver that links diverse metabolic pathways (Walker, 2017).

Lysine acetylation (Kac) is also induced under ketogenic conditions, as a substantial rise in acetyl-CoA feeds the production of β-OHB(Roberts et al., 2017; Sato et al., 2017). Thus, Kac occurs in parallel with Kbhb, as evident in liver whole-cell lysates from starved mice (Figure S3A). An important distinction to make is whether Kac and Kbhb target similar or distinct groups of proteins under these conditions. We thereby compared our Kbhb dataset with the previously determined lysine acetylome from 24 hr fasted mouse liver (Rardin et al., 2013b). Since this dataset was generated from mitochondrial fractions, we limited our analysis to mitochondrial localized proteins as determined by MitoMiner (Smith and Robinson, 2016). This analysis revealed that 82.8% of Kbhb-modified proteins and 72.4% of Kbhb-modified sites are also targeted by Kac (Figure 3F), indicating significant overlap of the pathways targeted by these two modifications.

### β-OHB regulates AHCY activity

One-carbon metabolism was among the top metabolic pathways enriched for Kbhb (Figure 3E). Notably, several enzymes implicated in methionine metabolism were identified by our MS analysis (Figure 4A). The importance of methionine metabolism for numerous cellular functions, ranging from biosynthesis of lipids and metabolites to epigenetic regulation through methylation of DNA, RNA and histones(Serefidou et al., 2019) (Haws et al., 2020) (Ducker and Rabinowitz, 2017), prompted us to determine the effects of Kbhb on this pathway. We focused on S-adenosyl-L-homocysteine hydrolase (AHCY), a rate-limiting enzyme that hydrolyzes S-adenosylhomocysteine (SAH) to homocysteine and adenosine (Figure 4A). Complete loss of AHCY leads to embryonic lethality(Dickinson et al., 2016), while AHCY deficiency in humans results in a number of pathological consequences including neurodevelopmental disorders, myopathy, hepatocellular carcinoma and early childhood death(Baric, 2009; Stender et al., 2015).

**Figure 4.**
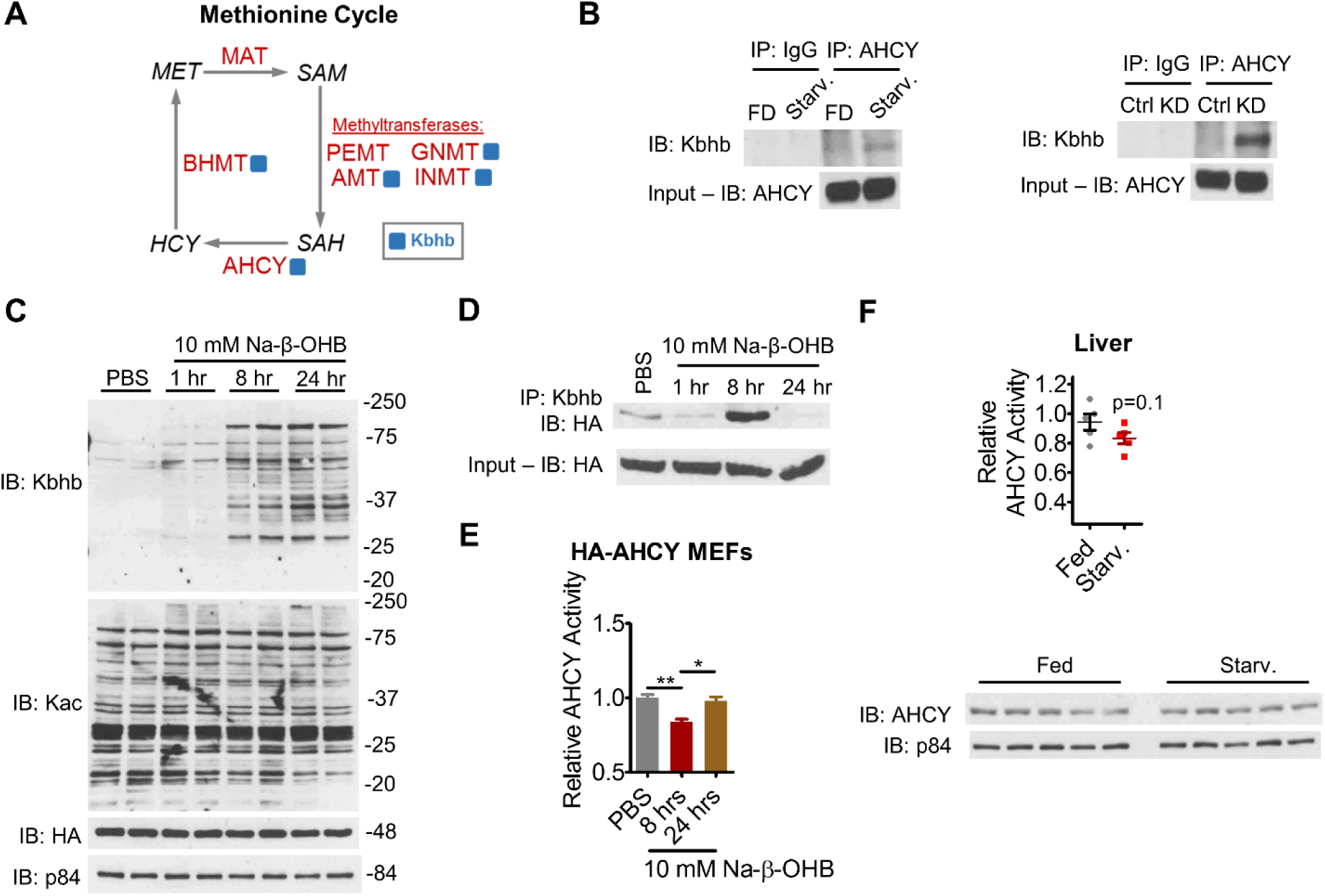
β-hydroxybutyrate inhibits AHCY enzymatic activity. (A) Simplified schematic of the methionine cycle. Enzymes are in red, metabolites are in black, and blue square indicates identification of Kbhb site. MAT – s-adenosylmethionine synthetase; BHMT – betaine-homocysteine methyltransferase; AHCY – adenosylhomocysteinase; PEMT – phosphatidylethanolamine N-methyltransferase; GNMT – glycine N-methyltransferase; INMT - indolethylamine N-methyltransferase; AMT – aminomethyltransferase. MET – methionine; SAM – s-adenosylmethionine; SAH – s-adenosylhomocysteine; HCY – homocysteine. (B) Immunoprecipitation of AHCY from liver whole cell lysates. FD – fed; Starv. – 48 hr fast; Ctrl – control diet; KD – ketogenic diet. (C) Time course of Kbhb induction in HA-AHCY MEF cells. (D) Time course of Kbhb of AHCY in HA-AHCY MEF cells revealed by immunoprecipitation. (E) AHCY activity was measured as the rate of adenosine production from SAH hydrolysis in HA-AHCY MEF whole cell lysates. One-way ANOVA, *=p<0.05, **=p<0.01. (F) AHCY was immunoprecipitated from liver whole cell lysates and its activity was measured as in (E), unpaired Student’s t-test. AHCY protein levels from liver whole cell lysates are shown below. Starv. – 48 hr fast.

Despite the biological importance of AHCY, little is known about its regulation. Thus, we sought to validate AHCY as a target of Kbhb by immunoprecipitation. Western blots show that both starvation and KD induce Kbhb on AHCY compared to control conditions (Figure 4B). To further confirm Kbhb of AHCY, we stably expressed HA-tagged AHCY in MEF cells. Comparable to WT MEFs, 10 mM Na-β-OHB treatment induced global protein Kbhb in a time-dependent manner in HA-AHCY MEFs (Figure 4C). Importantly, and in contrast to liver, sodium β-OHB treatment did not induce global protein acetylation (Figure 4C). Immunoprecipitation of HA-AHCY revealed a robust Kbhb signal for AHCY 8 hr after treatment that was attenuated by 24 hr (Figure 4D). These data show that AHCY is a bona fide target of Kbhb.

AHCY catalyzes the hydrolysis of SAH to homocysteine, freeing adenosine in the process. To determine if Kbhb of AHCY coincides with a change in its enzymatic activity, we measured the rate of adenosine production from SAH hydrolysis in HA-AHCY MEFs. While protein levels remained constant (Figure 4C), AHCY activity was inhibited at 8 hr following 10 mM Na-β-OHB treatment and returned to baseline by 24 hr, mirroring the transient Kbhb signal (Figure 4E). Similar timing of AHCY inhibition in response to 10 mM Na-β-OHB was observed in WT MEFs and HEK293T cells (Figure S4A-B). In starved liver, AHCY activity was attenuated while protein levels were unaltered (Figure 4F). Together, these results indicate that Kbhb inhibits AHCY activity.

### Kbhb of lysines within the NAD^+^ binding interface of AHCY inhibits its activity

Considering Kbhb of AHCY parallels inhibition of its enzymatic activity, we sought to gain insight into how Kbhb may elicit this regulation. To do so, we examined the crystal structure of AHCY in its NAD^+^-bound form (Figure 5A). AHCY functions as a tetramer, comprised of two homodimers of structurally identical ∼48-kDa subunits that each contain a substrate binding domain, cofactor binding domain and C-terminal loop (Grbesa et al., 2017). NAD^+^ binding to AHCY depends on the interaction between C-terminal loops of neighboring subunits, which together constitute one cofactor binding and substrate binding site (Wang et al., 2014). Four of the 6 Kbhb sites identified by our proteomics analysis are in close proximity to this critical interface (K188, K204, K389 and K405) while 2 are located on the surface, quite distally from the active site (K20 and K43). Our *in silico* analysis shows that K188 is functionally important for forming hydrogen bond interactions with the C-terminus of each homodimer and D223. K389 also participates in hydrogen bond interactions between S420 and D422 on the C-terminal loops, while the amine nitrogen of K405 interacts with the carboxyl group of E264. These interactions seemingly help support the NAD^+^ binding site conformation and, in turn, the nearby substrate binding site. Kbhb of these lysine residues would introduce a positive charge, predictive of disruption of these interactions and stability of this interface. Ultimately, an unfavorable conformational change could impair enzymatic activity.

**Figure 5.**
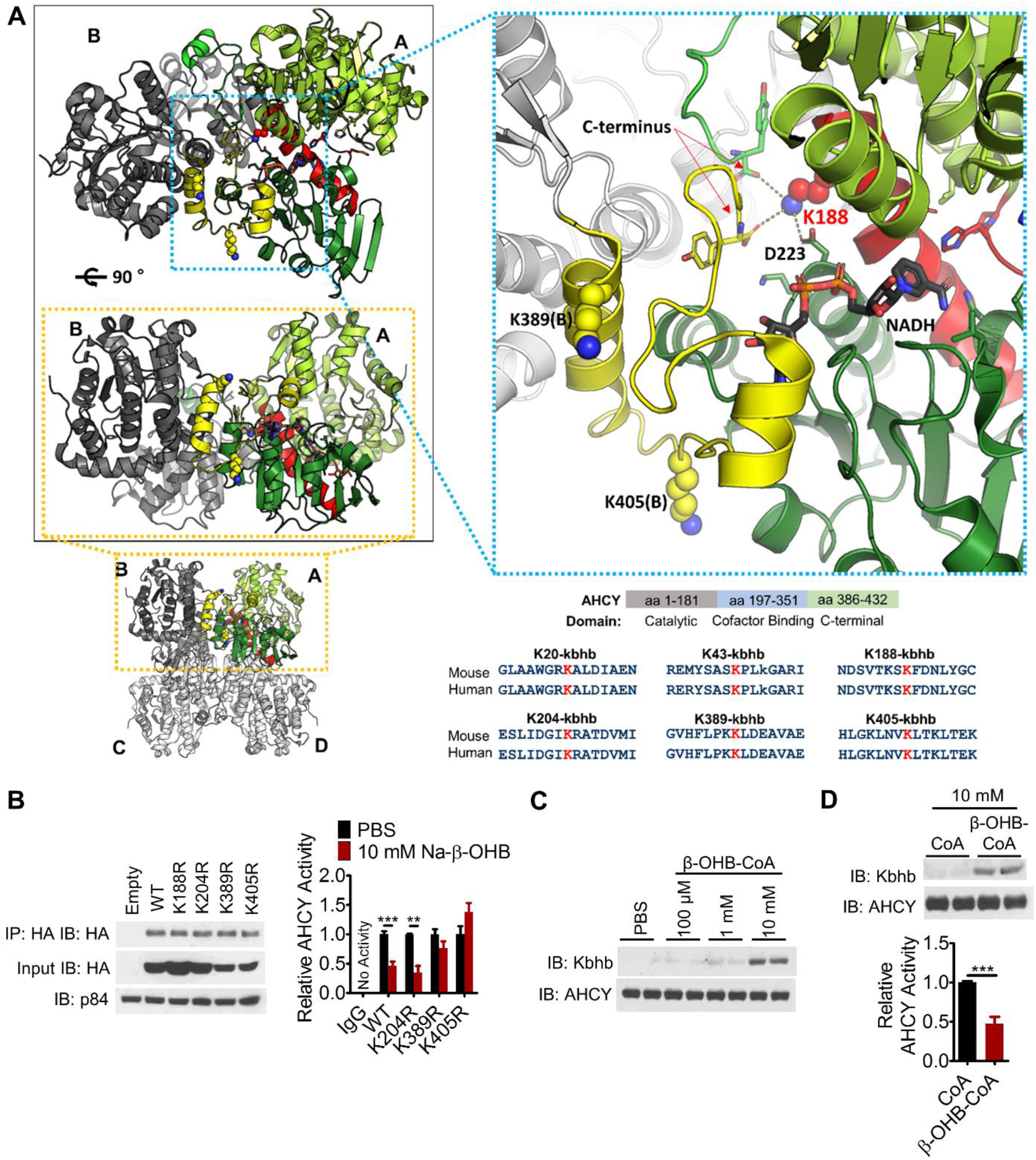
β-hydroxybutyrylation of K389 and K405 inhibits AHCY enzymatic activity. (A) Crystal structure of AHCY (PDB ID: 4YVF) bound to NAD^+^/NADH. Four identical monomers form 2 dimers (AB and CD) and, in turn, 1 tetramer. The substrate binding domain (light green) and cofactor binding domain (dark green) of A, and the C-terminal loop of B (yellow), constitute one active site. The connecting helix and loop between substrate binding and cofactor binding domains are shown in red. Kbhb sites are blue spheres (K188, K389[B], and K405[B]). The stick model of NAD^+^/NADH has black carbons. Dashed lines mark residues within hydrogen bond distance from K188. Structures were generated with PyMol; Bottom right – conservation of AHCY Kbhb sites from mouse to human. (B) HEK293T cells were transfected with WT or the indicated lysine mutant plasmid, treated for 10 hr with 10 mM sodium-β-hydroxybutyrate (Na-β-OHB) to induce Kbhb, then transfected AHCY was immunoprecipitated by its HA tag and assayed for enzymatic activity; values are normalized to PBS for each plasmid (see also Figure S5b). Unpaired Student’s t-test, **=p<0.01. ***=p<0.001. (C) Recombinant AHCY protein was incubated with increasing concentrations of β-hydroxybutyryl-CoA (β-OHB-CoA) for 2 hr at 37°C. (D) AHCY activity was measured from recombinant protein incubated with CoA or β-OHB-CoA as in (C). Unpaired Student’s t-test, ***=p<0.001.

To test our hypothesis, we mutated candidate lysine residues to arginine (K->R), a structurally similar amino acid with the same charge, to mimic an unmodified lysine that cannot be subsequently β-hydroxybutyrylated. HEK293T cells, whose response to 10 mM Na-β-OHB treatment is comparable to HA-MEFs (Figure S5A), were transfected with WT, K188R, K204R, K389R or K405R HA-AHCY plasmids and treated with 10 mM Na-β-OHB for 10 hr to induce Kbhb. To avoid influence from endogenous AHCY, we immunoprecipitated the transfected AHCY by its HA tag and directly measured enzymatic activity. All constructs were expressed and immunoprecipitated at comparable levels (Figure 5B). Of the four mutations, only K188R had a substantial effect on its own, almost completely blocking AHCY activity, demonstrating that K188 mediates a vital structural and functional interaction (Figure S5B). Next we analyzed the contribution of each residue under Kbhb. Compared to PBS treatment, 10 mM Na-β-OHB inhibited WT AHCY activity (Figure 5B). The K204R mutant was inhibited by β-OHB equally to WT, demonstrating a negligible effect of Kbhb at this site (Figure 5B). In contrast, K389R and K405R abolished β-OHB-induced inhibition of AHCY activity (Figure 5B). Considering the position of these residues and the crystal structure modeling, we posit that Kbhb disrupts the critical C-terminal loop interactions involving K389 and K405, destabilizing the NAD+ binding site conformation and impairing enzymatic activity.

A synergistic effect of Kbhb at several sites including K389, K405 and perhaps others is likely, ultimately acting together in altering AHCY enzymatic function. Thus, we next determined the direct effect of β-hydroxybutyrylation on AHCY activity in a cell free assay. Recombinant AHCY protein was incubated with β-hydroxybutyryl-CoA for 2 hr at 37°C and formation of Kbhb was monitored by western blot. β-hydroxybutyryl-CoA induced Kbhb on AHCY in a concentration-dependent manner, in the absence of any potential acylating enyzme (Figure 5C). Compared to incubation with CoA, β-hydroxybutyryl-CoA reduced AHCY enzymatic activity (Figure 5D), similarly to the effect of β-OHB observed in cells. These data show that Kbhb directly inhibits AHCY activity.

## DISCUSSION

β-OHB is a dynamic metabolite implicated in signaling and epigenetic regulation. Here we report an unappreciated role for β-OHB in the regulation of hepatic metabolism. Indeed, using high-throughput proteomic analysis of lysine β-hydroxybutyrylation in liver we reveal Kbhb as a widespread posttranslational modification of non-histone proteins (Figure 6). Focusing on the methionine cycle enzyme AHCY, we demonstrate the ability of Kbhb to alter the enzymatic activity of a key metabolic regulator. Finally, we implicate global protein Kbhb broadly in biological contexts by illuminating its presence upon dietary, disease and potentially therapeutic interventions.

**Figure 6.**
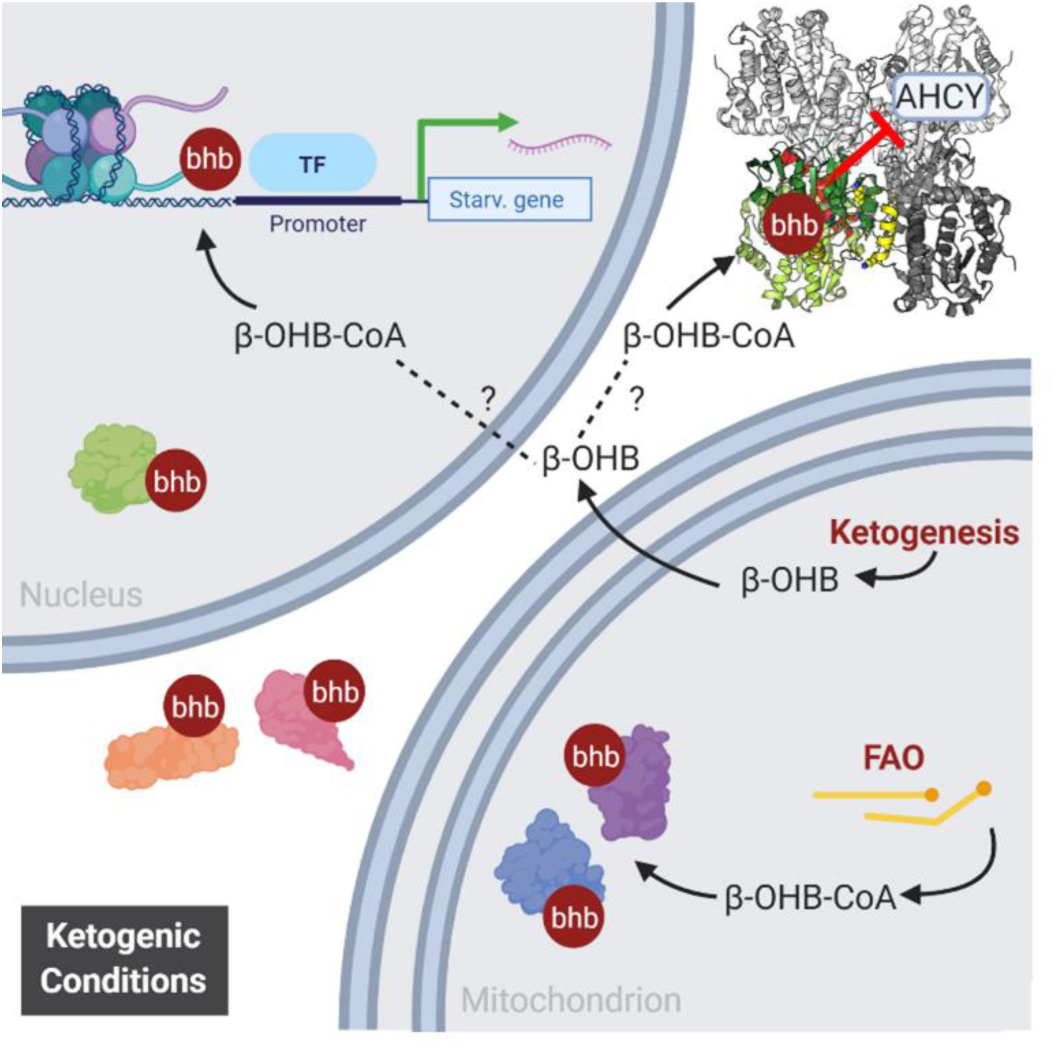
Model of the Link Between Ketogenic Metabolism and Global Protein Lysine β-hydroxybutyrylation.

Interestingly, the induction of global protein Kbhb appears highly tissue-specific to the liver and kidney. During ketogenesis, the liver produces most, if not all, β-OHB and its own concentration parallels that of the blood(Puchalska and Crawford, 2017; Tognini et al., 2017). Reabsorption of β-OHB by the kidney is proportionate to blood levels, and kidney β-OHB levels are markedly increased by a 24 hr fast(Puchalska and Crawford, 2017). Under these conditions the metabolic tasks of the liver and kidney share some similarities in that both organs help maintain circulating glucose by gluconeogenesis and provide ketone bodies through ketogenesis (although renal ketogenesis is still debated) (Puchalska and Crawford, 2017). While hepatic and renal β-OHB levels rise, it flows down its concentration gradient through solute carrier 16A 1 and 7 (MCT1 and MCT2) into extrahepatic tissues where it is rapidly metabolized to acetyl-CoA for terminal oxidation via the TCA cycle. Since heavy oxidizers of β-OHB such as the heart, skeletal muscle and brain do not display Kbhb induction (Figures 1 and 2), it appears that Kbhb is tied to β-OHB levels in tissue. This notion is in keeping with the dose-dependent induction of Kbhb *in vitro* and *in vivo* by β-OHB. Notably, Kbhb of p53 was recently reported in the thymus (Liu et al., 2019), a typically less metabolic tissue, therefore indicating that Kbhb on specific proteins may be more nuanced from tissue to tissue.

Ultimately, it is the activated, CoA-bound form of β-OHB that serves as the substrate for Kbhb(Xie et al., 2016), however the exact source of acylation-promoting acyl-CoAs for many PTMs including β-OHB is unclear. There are several possibilities for β-OHB-CoA. Acyl-CoA synthetase short chain 2 (ACSS2) is a nucleo-cytoplasmic enzyme that converts acetate to acetyl-CoA, supporting acetylation of histones (Mews et al., 2017) (Sahar et al., 2014), as well as crotonate to crotonyl-CoA and histone crotonylation (Sabari et al., 2018). It is theorized that ACSS2 acts broadly to generate activated CoA forms of other short chain fatty acids and β-OHB is a candidate substrate(Sabari et al., 2017). Another possibility is that β-OHB-CoA is generated by flux through mitochondrial pathways as an intermediate metabolite(Ringel et al., 2018). Two forms of β-OHB-CoA are known to be generated through fatty acid β-oxidation; acetoacetyl-CoA is converted to *L*-β-OHB-CoA by mitochondrial 3-hydroxyacyl-CoA dehydrogenase (Reed and Ozand, 1980), and a yet to be defined enzyme generates *S*-β-OHB-CoA from an acyl-CoA precursor (Robinson and Williamson, 1980). β-oxidation is ramped up under fasting, starvation and ketogenic diet, and once rates approach saturation, β-OHB-CoA concentration could rise substantially to out-compete other acyl-CoA species for reactivity with lysines or as a substrate for a yet to be defined enzymatic writer. Our *in vitro* results demonstrate non-enzymatic Kbhb at high, yet physiologically relevant concentrations of β-OHB-CoA(Wagner et al., 2017). Nevertheless, an enzyme-catalyzed mechanism is also likely for Kbhb deposition. The acetyltransferase p300 binds an array of acyl-CoAs(Kaczmarska et al., 2017), writes several other acyl PTMs(Huang et al., 2018a; Huang et al., 2018b; Sabari et al., 2018) and has been shown to facilitate histone Kbhb in a cell-free assay, hence it is a candidate β-hydroxybutyrylating enzyme.

Similar to what is observed for other acylations (Wagner and Hirschey, 2014) (Choudhary et al., 2009), several macronutrient pathways were enriched for Kbhb including fatty acid β-oxidation, TCA cycle, glycolysis, amino acid metabolism, urea cycle and ketogenesis. Several rate-limiting enzymes therein have more than 10 Kbhb sites: CPS1 (urea cycle); CS (TCA cycle); HMGCS2 (ketogenesis). These metabolic networks are paramount for the fuel switching that must occur in the liver for a proper adaptation to ketogenic conditions, whereby fatty acid oxidation and amino acid catabolism fuel ketogenesis while gluconeogenesis maintains circulating glucose levels. Therefore, Kbhb appears to link cellular metabolism to the regulation of vital pathways at global proteomic level. Intriguingly all the enzymes participating in the production of β-OHB are modified by Kbhb. These include the rate-limiting HMGCS2 and the mitochondrial 3-hydroxyacyl-CoA dehydrogenase (HADH) which produces β-OHB-CoA(Reed and Ozand, 1980). Thus, we envision that Kbhb feeds back onto the pathways that generate its parent molecule as well as its activated CoA form.

Protein acetylation, which is also induced by fasting and a ketogenic diet (Zhao et al., 2010) (Roberts et al., 2017), occurs in parallel with Kbhb. When limiting analysis to mitochondria, ∼75% of the proteins and sites of Kbhb overlap with Kac (Rardin et al., 2013b). Since our identification of Kbhb peptides by LC-MS/MS was performed in the presence of increased acetylation, it is possible that β-OHB-CoA competes with acetyl-CoA for these sites. Maximal ketogenic conditions might favor higher concentrations of β-OHB-CoA, since to a significant degree the fate of acetyl-CoA in the liver is β-OHB (Newman and Verdin, 2017). However, a limitation of our MS data is that the relative stoichiometry of Kbhb to Kac or unmodified lysine is not defined. The stoichiometry at specific lysine residues may prove crucial for the regulation of metabolic enzymes, as hypothesized for the competition between other acyl lysine modifications such as acetylation, malonylation and succinylation (Nishida et al., 2015).

Interestingly, the liver and kidney share 1-carbon (1C) metabolism, a metabolic network that integrates several nutrients to fuel a variety of physiological processes, including nucleotide metabolism, amino acid homeostasis, lipid biosynthesis, redox balance and methylation metabolism(Ducker and Rabinowitz, 2017). Several enzymes of the methionine cycle, a major arm of 1C metabolism, were β-hydroxybutyrylated in liver (Figure 3). The methionine cycle is of particular physiological importance because it regulates all cellular trans-methylation reactions by controlling the ratio of S-adenosylmethionine (SAM) and S-adenosylhomocysteine (SAH), and is therefore a critically controlled metabolic pathway. Notably, several enzymes identified in our (LC)-MS/MS analysis belong to this pathway. These include serine hydroxymethyltransferase (SHMT1), betaine-homocysteine methyltransferase (BHMT), glycine-N-methyltransferase (GNMT) and S-adenosyl-L-homocysteine hydrolase (AHCY) (Figure 4a). These enzymes function at several key steps of the cycle and together contribute to a tight control of SAM and SAH cellular levels(Girgis et al., 1998; Pajares and Perez-Sala, 2006; Yeo and Wagner, 1992), suggesting that protein β-hydroxybutyrylation may play a role in methionine homeostasis under conditions of metabolic stress, such as prolonged fasting.

Many of the Kbhb sites identified in this study overlap with lysine residues important for enzymatic function, giving Kbhb the potential to modulate enzymatic activity. For example, carbamoyl phosphate 1 (CPS1), a rate-limiting enzyme in the urea cycle, is modified at 35 sites, including lysine residues previously linked to posttranslational control of its activity(Tan et al., 2014). This is the case of AHCY, whose β-hydroxybutyrylation decreases its enzymatic activity, demonstrating a functional relevance of the modification. In addition to the sites directly affecting activity, AHCY was modified at two additional sites in the N-terminal domain, which we hypothesize could affect protein-protein interactions or subcellular localization of AHCY(Grbesa et al., 2017). Future studies will help define the full array of functional consequences of Kbhb on target proteins and whether, as observed for AHCY, Kbhb targets other metabolic enzymes in a similar manner.

Global protein β-hydroxybutyrylation could play a significant role in metabolic adaptation in certain diseases. The ketogenic pathway has been linked to cancer (Kang et al., 2015; Poff et al., 2014), for example a ketogenic diet can elicit poorly understood anti-tumor effects(Weber et al., 2018). Since cancer cells often have impaired mitochondrial function, they are unable to utilize ketone bodies as an energy source(Maurer et al., 2011). This opens the possibility that, when held under a ketogenic diet, cancer cells may accumulate β-OHB and consequently β-OHB-CoA and Kbhb. Cancer cells are highly dependent on methionine metabolism and several studies report AHCY as a possible therapeutic target(Beluzic et al., 2018; Chayka et al., 2015). Moreover, in keeping with our results, variations in methionine metabolites have been linked to beneficial effects of ketogenic diets(Pissios et al., 2013), which presumably operate, at least in part, through protein Kbhb.

## AUTHOR CONTRIBUTIONS

K.B.K., C.M.G., H.H., Y.Z., F.Q., C.L.P. and P.S.-C. designed the study. K.B.K., C.M.G., H.H., J.F. and P.C. conducted experiments and collected data. J.K. and F.Q. performed bioinformatic analyses. K.B.K., C.M.G. and P.S.-C. wrote the paper, with input from all authors.

## ACKNOWLEDGEMENTS

We thank all members of the Sassone-Corsi lab for discussions and support. P.S.-C. lab is supported by NIH grants R21DK114652, R21AG053592 and a Challenge Grant of the Novo Nordisk Foundation (NNF-202585). K.B.K was supported by NIH-NINDS, T32 5T32NS045540 and currently F32DK121425. C.M.G was supported by the National Cancer Institute of the US National Institutes of Health (NIH T32 2T32CA009054-36A1) and by European Research Council (ERC MSCA-IF-2016 MetEpiClock 749869). Y.Z. was supported by NIH grants R01GM115961, R01DK107868, and R01DK118266.

## ACCESSION NUMBERS

The mass spectrometry proteomics data have been deposited to the ProteomeXchange Consortium via the PRIDE (Vizcaino et al., 2016) partner repository with the dataset identifier PXD016654 (Reviewer account details: Username; Password, UEbpEtS5). Data are also available in Supplemental Table S1.

## MATERIALS AND METHODS

### Animals and Interventions

Animal experiments were conducted in accordance with the guidelines of the Institution Animal Care and Use Committee (IACUC) at UC Irvine. Consideration was given to the ARRIVE guidelines during the development of the experimental design. Unless otherwise stated, adult male (20-30 g) C57BL/6 mice were group housed under a 12 hr light, 12 hr dark cycle and fed *ab libitum* with normal chow (2020X Envigo, Teklad). For fasting experiments, food was withdrawn at *Zeitgeber* time (ZT) 8, and tissue was harvested at ZT8 either 24 or 48 hr later. Caloric restriction samples were collected as described in Sato et al. Cell 2017. Briefly, 19-29 week-old mice were fed either control chow (Harlan TD.120685) or 30% calorie restricted chow (Harlan TD.120686) for 25 weeks and liver samples (examined in the present study) were harvested at ZT8. Tissues from mice fed control chow (4 weeks, TD.150345 Envigo, Teklad), ketogenic diet (4 weeks, TD.160153 Envigo, Teklad) or 10% w/w 1,3-butanediol diet (3 weeks, TD. 160257 Envigo, Teklad) were also harvested at ZT8. To induce type I diabetes mellitus (TIDM), mice were injected IP with 200 mg/kg streptozotocin (Caymen Chemical, 13104). Blood measurements and tissue harvest occurred 4 days after injections when the induction of TIDM was evident.

### Immunoprecipitation

Whole cell lysates were prepared as described in the western blot section. 500 µg of protein was incubated overnight at 4°C with primary antibody as indicated (anti-AHCY [MBL RN126PW], anti-HA [Millipore 05-904], anti-IgG Rb (Invitrogen 10500C). 20 µl of Protein G Dynabeads (Thermo Fisher Scientific) were added to the samples and incubated on a rocker at 4°C for 2 hr. Beads were then washed 4X with RIPA buffer (see western blot section). For western blot, bound proteins were eluted directly in SDS loading buffer. For AHCY activity, beads were added directly to the reaction mix. For primary antibodies raised in rabbit, an anti-light chain, HRP-conjugated secondary antibody (Millipore MAB201P) was used to develop blots.

### Subcellular Fractionation

Mouse livers were homogenized with a motorized tissue grinder in STM buffer (Tris-HCl pH 7.4, 5 mM MgCl_2_, 250 mM Sucrose, 1 mM DTT, supplemented with 500 µM PMSF, Protease Inhibitor Cocktail [Roche, Basel, Switzerland], 10 mM nicotinamide and 330 nM TSA). Samples were then passed through a 100 µM filter, let on ice for 10 min and vortexed at max speed for 10 sec. Nuclei were pelleted by centrifugation at 800 g for 10 min at 4°C and set aside. The same centrifugation was repeated on the supernatant to further remove nuclei. Next, mitochondria were pelleted by centrifugation at 11,000 g for 10 min at 4°C and set aside. Again, centrifugation was repeated on the supernatant to further remove mitochondria. The supernatant, which we took as the cytosolic fraction, was finally centrifuged at max speed for 10 min to remove any remaining nuclear or mitochondrial contamination. The nuclear pellet was next resuspended in CYTO buffer (10 mM HEPES-NaOH pH 8.0, 25 mM KCl, 650 µM spermidine, 1 mM EDTA, 1 mM EGTA, 340 mM sucrose, 1% NP40, 1 mM DTT, supplemented with 500 µM PMSF, Protease Inhibitor Cocktail, 10 mM nicotinamide and 330 nM TSA) and centrifuged at 800 g for 10 min at 4°C to clean the fraction. The final nuclear pellet was lysed in RIPA buffer (50 mM tris-HCl pH 8.0, 150 mM NaCl, 5 mM EDTA, 15 mM MgCl^2^ and 1% NP-40, supplemented with 500 µM PMSF, Protease Inhibitor Cocktail, 10 mM nicotinamide and 330 nM TSA). The Mitochondrial pellet was resuspended in STM buffer and centrifuged at 11,000 g for 10 min at 4°C to clean the fraction. The final mitochondrial pellet was lysed in RIPA buffer. Both nuclear and mitochondrial fractions were then sonicated at 60% amplitude, 5 sec on 5 sec off, for 20 sec. Samples were then centrifuged at max speed for 10 min at 4°C to remove debris and membranes. From these fractions, samples were prepared for western blotting as described in the western blot methods section.

### Histone Extraction

Liver tissue was homogenized with a motorized tissue grinder in extraction buffer (PBS, 0.5% triton X 100 [v/v], supplemented with 500 µM PMSF, Protease Inhibitor Cocktail, 10 mM nicotinamide and 330 nM TSA) and lysed on ice for 10 min. Samples were centrifuged at 6,500 g for 10 min at 4°C to pellet nuclei, which were then washed in extraction buffer by repeating the previous centrifugation. Nuclei were resuspended in 0.2 M HCl overnight at 4°C. Following centrifugation at 6,500 g for 10 min at 4°C, the supernatant was mixed with a final concentration of 33% TCA to precipitate histones on ice for 30 min. Histones were pelleted by centrifugation at max speed for 10 min at 4°C. Pellets were then washed with ice cold 100% acetone and centrifugation was repeated for 2 washes. Final histone pellets were air dried for 20 min at room temperature, dissolved in ddH_2_O and prepared with SDS loading buffer for western blot as described. The amount of histones per sample was compared by ponceau staining and normalized accordingly for subsequent blotting.

### β-hydroxybutyrate and Glucose Measurements

Blood β-hydroxybutyrate levels were measuring with a Nova Max Plus™ β-ketone meter (Nova Biomedical). Blood glucose was measured with a Contour blood glucose monitor (Bayer). Readings were acquired by a ∼2 mm tail cut.

### AHCY Activity Assay

To measure the enzymatic activity of AHCY from total lysates, samples were incubated in homogenization buffer, provided in the Adenosylhomocysteinase (AHCY) Activity Fluorometric Assay Kit (Biovision, K807-100), and let on ice for 15 min at 4°C, followed by centrifugation at 14,000 rpm for 10 min at 4°C. Pellets were then resuspended in AHCY assay buffer. Activity of AHCY from total lysates and immunoprecipitated AHCY was measured according to the manufacturer’s instructions. The assay was performed in 96-well white plates and fluorescence was measured in kinetic mode for 30-45 min at 37°C using a Varioskan LUX multimode microplate reader (Thermo Fisher Scientific).

### AHCY Mutagenesis

HA-AHCY was purchased from VectorBuilder and mutations were introduced by PCR-based mutagenesis using Q5 Site-Directed Mutagenesis Kit (NEB) according to manufacturer’s instructions. Primers were as follows: K405R FOR 5’-acctgggcaagctgaatgtgaggctgaccaagc-3’, REV 5’-gcttggtcagcctcacattcagcttgcccaggt-3’; K389R FOR 5’-gggttcacttcctgcctaagaggctggatgagg-3’, REV 5’-cctcatccagcctcttaggcaggaagtgaaccc-3’; K188R FOR 5’-attctgtcaccaagagcaggtttgacaacctctatgg-3’, REV 5’-ccatagaggttgtcaaacctgctcttggtgacagaat-3’; K204R FOR 5’-gatggcatcagacgggccaca-3’ REV 5’-tatgagggactcccggca-3’. Mutagenesis was confirmed by sequencing.

### AHCY Protein Expression and Purification

Mouse AHCY residues 1-432 were cloned into the pHIS-parallel vector, which adds a His_6_ tag at the N-terminus, using the BamHI and Sal1 sites. His_6_-AHCY was expressed in the BL21 (DE3) strain of *E. coli*. Cells were grown at 37° C with shaking until they reached an OD_600_ of 0.8, whereupon His_6_-AHCY expression was induced with 0.5 mM isopropyl-β-D-thiogalactopyranoside (IPTG) and then grown for an additional 16 h at 18° C. Cells were harvested by centrifugation at 4000 x rpm and resuspended in lysis buffer (50 mM Tris pH 8.0, 500 mM NaCl and 20 mM imidazole). Cells were lysed using a microfluidizer followed by sonication on ice (15 seconds on, 45 seconds off for four pulses at 40% amplitude with a 1/4” microtip). His_6_-AHCY was isolated by Ni^2+^-nitrilotriacetic acid (Ni-NTA) affinity chromatography (Qiagen) using standard procedures and eluted with 50 mM Tris buffer pH 8.0, 500 mM NaCl and 300 mM imidazole. The His_6_ tag was cleaved with His_6_-TEV protease overnight at 4° C. The Ni-NTA eluate was then diluted with lysis buffer to lower the concentration of imidazole to 40 mM and subsequent Ni-NTA affinity chromatography was performed to remove His_6_ tag and His_6_-Tev.

Preparation of NAD+-bound holoenzyme was performed as previously described (Manszewski et al., 2017). Briefly, 5 mL of apo AHCY (at a concentration of 12 mg/mL) was mixed with 10 mL of saturated ammonium sulfate solution at pH 3.3 and incubated on ice for 10 minutes. The protein was then pelleted by centrifugation at 10,000 x rpm for 15 minutes at 4° C. The precipitate was dissolved in 5 mL of lysis buffer and mixed with 10 mL of saturated ammonium sulfate solution at pH 3.3, incubated and centrifuged again. Supernatant was discarded and the pellet was again diluted in lysis buffer and mixed with 10 mL of saturated ammonium sulfate solution at pH 7.0. The mixture was centrifuged again and the resulting pellet resuspended in lysis buffer. Protein was incubated overnight at 4° C with 12-fold excess of NAD+. The preparation was further purified by Superdex 200 gel filtration chromatography (GE Healthcare) into 20 mM Tris pH 8.0, 50 mM NaCl. The protein was aliquoted, frozen in liquid nitrogen, and stored at −80° C for storage.

### AHCY *in Vitro* Acylation

A cell free assay to induce β-hydroxybutyrylation of AHCY was performed as described previously with adjustments (Kaczmarska et al., 2017). Reactions were set up in reaction buffer (25 mM Tris-HCl pH 8.0, 100 mM NaCl, 1 mM DTT, 100 µM EDTA, 10% glycerol, supplemented with Protease Inhibitor Cocktail, 10 mM nicotinamide and 100 ng/ml TSA), with 2 µg AHCY and 100 µM, 1 mM or 10 mM DL-β-hydroxybutyryl-CoA lithium salt (Sigma H0261), for 2 hr in a 37°C water bath. Reactions were either stopped with addition of SDS loading buffer for western blot or loaded directly into AHCY activity assay wells for measurement.

### Cell Culture

Mouse embryonic fibroblasts (MEF), human embryonic kidney (HEK) 293T and mouse hepatoma (Hepa1c1c) cells were cultured in DMEM (4.5 g l-^1^ glucose, Hyclone) supplemented with 10% FBS (Gibco) and 100 units (penicillin), 100 µg (streptomycin) antibiotics (Gibco). To generate stable expression of HA-AHCY in MEF cells, lentiviral HA-AHCY expression vector (VectorBuilder) was transfected into HEK293T cells together with psPAX2 and VSVG second-generation lentiviral packaging systems using BioT reagent (Bioland Scientific LLC), according to manufacturer’s instructions. After 48h, lentiviral particles in the medium were collected, filtered and used to infect MEFs. 48h post transduction, the cells were subjected to hygromycin (Sigma) selection. Overexpression efficiency was verified by western blotting. (R)-(-)-3-hydroxybutyric acid sodium salt (Sigma 298360) was added to the culture media at the indicated concentrations. For dose response experiments, cells were treated for 24 hr. For time course experiments, treatment times were as indicated on figures and in legends. 293T were transfected with WT or mutant HA-AHCY plasmids using BioT reagent (Bioland Scientific LLC), according to manufacturer’s instructions.

### Western Blot

Whole cell lysates were prepared by homogenizing cells or tissue with a motorized tissue grinder in RIPA buffer (see subcellular fractionation section for recipe). Samples were lysed on ice for 20 min, vortexed briefly and sonicated at 60% amplitude, 5 sec on 5 sec off, for 4 cycles. Membranes and debris were removed by centrifugation at max speed for 10 min at 4°C. Protein concentration was measured via the Braford method with Protein Assay Dye from BioRad (Hercules, CA). Ten to 50 µg protein was loaded into a 12% SDS-PAGE gel. Following transferal to a nitrocellulose membrane (Thermo Fisher Scientific, Waltham, MA), we blocked with 5% milk in TBST (0.1% Tween-20, Tris Buffered Saline) for 2 hr at room temperature. Primary antibodies (Anti-3-hydroxybutyryllysine [PTM Biolabs 1201], anti-p84 [GeneTex GTX70220], anti-α-Tubulin [Sigma T5168], anti-COXIV [Cell Signaling 11967S], anti-acetyllysine [Cell Signaling 9441L], anti-AHCY [Abcam ab134966], anti-HA [Millipore 05-904]) were diluted in 5% milk TBST and incubated with membranes overnight at 4°C. Membranes were then incubated with HRP-conjugated secondary antibodies (Mouse IgG-HRP conjugate [EMD Millipore AP160P], Rabbit IgG-HRP linked [EMD Millipore 12-348]) for 1 hr at room temperature. Blots were developed with Immobilon Western Chemiluminescent HRP Substrate (Millipore, Burlington, MA) and HyBlot CL autoradiography film (Denville Scientific, Holliston, MA).

### Proteomics

Livers from starved (48 hr fasted) mice were cut into small pieces, washed with cold PBS and homogenized on ice in lysis buffer (8 M urea, 2 mM EDTA, 3 μM TSA, 50 mM NAM, 5 mM DTT and 1% Protease Inhibitor Cocktail III). The remaining debris was removed by centrifugation at 18,000 × g at 4 °C for 3 min. Proteins in the cell lysate were reduced with 10 mM DTT for 1 hr at 37 °C, alkylated with 20 mM iodoacetamide for 45 min at room temperature in darkness, and the excess iodoacetamide was blocked by 20 mM cysteine. Then the protein sample was diluted by adding 100 mM NH_4_HCO_3_ to reduce the urea concentration to 1 M. Trypsin was added at 1:50 trypsin-to-protein mass ratio for the first digestion overnight and 1:100 trypsin-to-protein mass ratio for a second 4 hr digestion. Finally, 18 mg of protein was digested for subsequent experiments. Peptides were fractionated into 23 fractions by high pH reverse-phase HPLC using an Agilent 300 Extend C18 column (5 μm particles, 4.6 mm ID, 250 mm length). Briefly, peptides were first separated using a gradient of 2%-60% acetonitrile in 10 mM ammonium bicarbonate at pH 10 over 90 min into 90 fractions. The peptides were then combined into 23 fractions and dried by vacuum centrifugation. To enrich Kbhb peptides, total peptides dissolved in NETN buffer (100 mM NaCl, 1 mM EDTA, 50 mM Tris-HCl, 0.5% NP-40, pH 8.0) were incubated with pre-washed pan anti-Kbhb beads (PTM Biolabs Inc., Chicago, IL) at 4 °C overnight with gentle shaking. The beads were washed four times with NETN buffer and twice with ddH_2_O. The bound peptides were eluted from the beads with 0.1% trifluoroacetic acid, and the eluted fractions were combined and vacuum-dried. Samples were dissolved in 0.1% formic acid and loaded onto a home-made capillary column (10 cm length with 75 um inner diameter) packed with Reprosil 100 C18 resin (3 µm particle size, Dr. Maisch GmbH, Ammerbuch, Germany). Peptide separation was performed with a gradient of 5-80% HPLC buffer B (0.1% formic acid in 90% acetonitrile, v/v) in buffer A (0.1% formic acid in water, v/v) at a flow rate of 300 nl/min over 60 min on an Eksigent UPLC system. The samples were analyzed by Orbitrap Velos Mass Spectrometers (ThermoFisher Scientific). A data-dependent procedure that alternated between one full mass scan followed by the top 20 most intense precursor ions was applied with 90 sec dynamic exclusion. Intact peptides were detected with a resolution of 30,000 at 400 m/z, and ion fragments were detected at 35%.

### Proteomic Bioinformatic Analysis

Database searching was performed with MaxQuant 1.3.0.5 (Cox and Mann, 2008) against the mouse proteome downloaded from UniProt. Trypsin was specified as cleavage enzyme allowing a maximum of 2 missing cleavages. Cysteine carbamidomethylation was set as a fixed modification. Methionine oxidation, N-terminal acetylation of proteins, and lysine β-hydroxybutyrylation were set as variable modifications. The false discovery rate (FDR) was set to 1% for peptide, protein and modification sites. Peptides identified from reverse or contaminant protein sequences, peptides with score below 40 or site localization probability below 0.75 were removed.The flanking sequence motif of lysine β-hydroxybutyrylation substrates were tested against all mouse background sequences with iceLogo (Colaert et al., 2009) using a p-value of 0.05.

### Statistical Analysis

Details of statistical analyses are indicated in figure legends and corresponding methods sections.

## SUPPLEMENTAL INFORMATION

### SUPPLEMENTAL FIGURE LEGENDS

**Figure S1.**
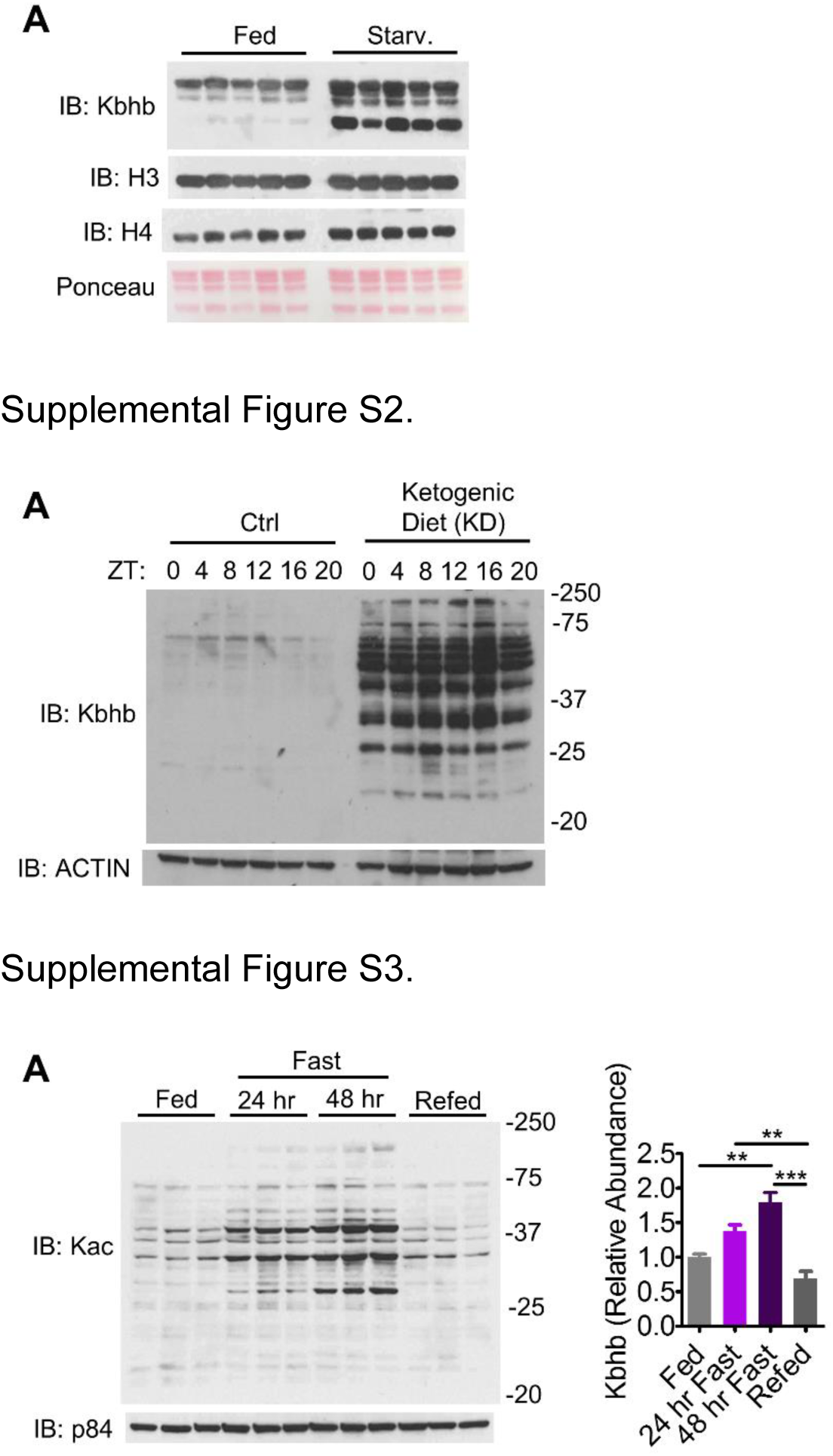
Related to Figure 1. (A) Western blot of histone extracts from fed and starved (48 hr fast) livers.

**Figure S2. Related to Figure 2**. (A) Whole cell lysates of livers harvested at 6 different time points over the circadian cycle. ZT0=lights on, ZT12=lights off.

**Figure S3. Related to Figure 3**. (A) Whole cell lysates from fed and starv. (48 hr fast) liver. n=3 replicates are quantified to the right, unpaired Student’s t-test, **=p<0.01, ***=p<0.001.

**Figure S4.**
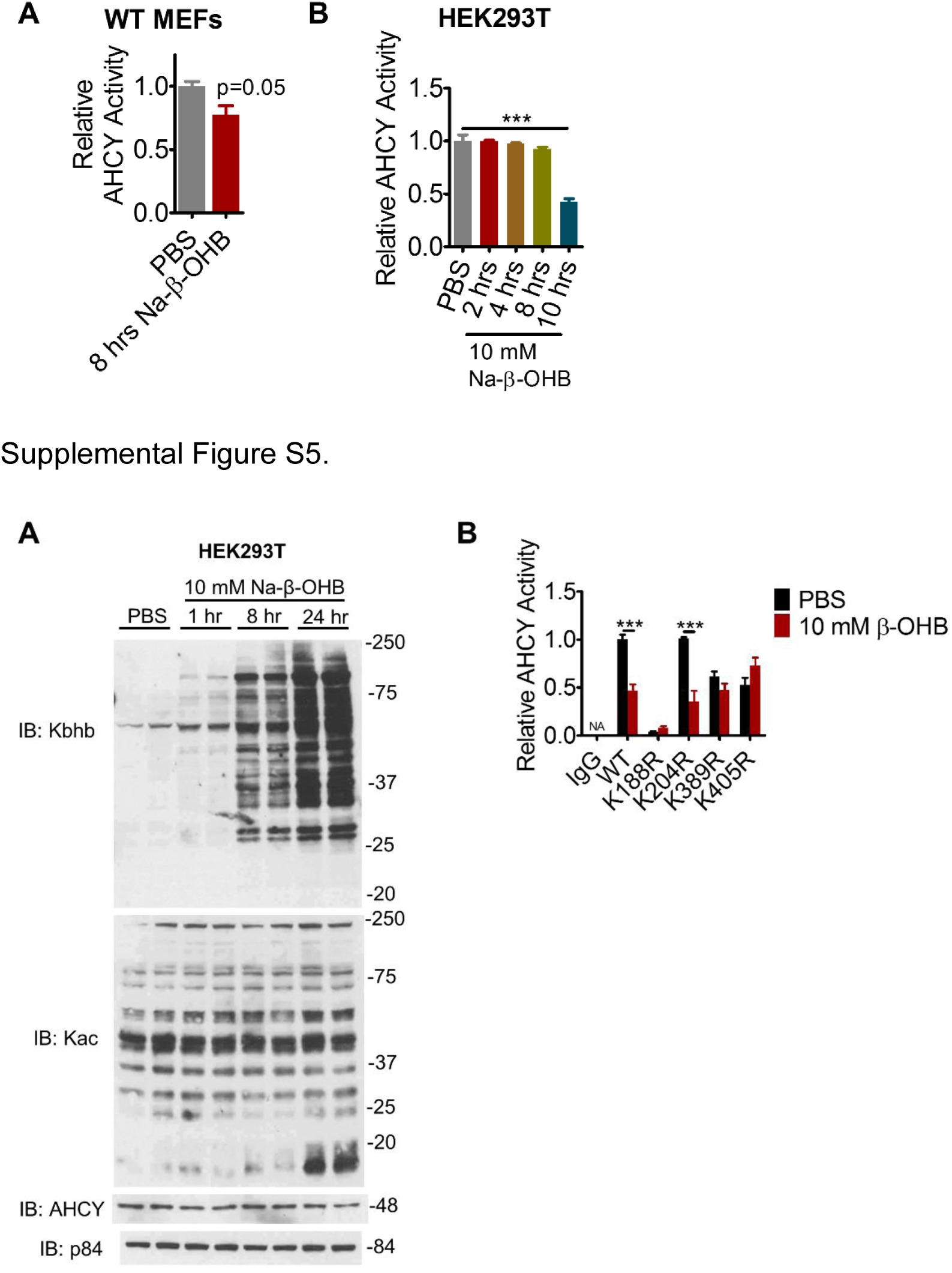
Related to Figure 4. (A-B) AHCY activity measured in whole cell lysates of WT MEF (A) and HEK293T (B) cells. a, unpaired Student’s t-test, n=3. (B) One-way ANOVA, ***=p<0.001, n=3-6.

**Figure S5. Related to Figure 5**. (A) Time course of Kbhb induction in HEK293T cells following sodium-β-hydroxybutyrate (Na-β-OHB) treatment. (B) HEK293T cells were transfected with WT or the indicated lysine mutant plasmid, treated for 8-10 hr with 10 mM sodium-β-hydroxybutyrate (Na-β-OHB) to induce Kbhb, then transfected AHCY was immunoprecipitated by its HA tag and assayed for enzymatic activity (as in Figure 5B). Values are normalized to WT PBS treated. T-test, ***=p<0.001.

**Table S1. Related to Figure 3, Identification of Kbhb sites in starved liver**.

## REFERENCES

Balasse, E.O., and Fery, F. (1989). Ketone body production and disposal: effects of fasting, diabetes, and exercise. Diabetes Metab Rev 5, 247–270.

Baric, I. (2009). Inherited disorders in the conversion of methionine to homocysteine. J Inherit Metab Dis 32, 459–471.

Beluzic, L., Grbesa, I., Beluzic, R., Park, J.H., Kong, H.K., Kopjar, N., Espadas, G., Sabido, E., Lepur, A., Rokic, F., et al. (2018). Knock-down of AHCY and depletion of adenosine induces DNA damage and cell cycle arrest. Sci Rep 8, 14012.

Cahill, G.F., Jr. (2006). Fuel metabolism in starvation. Annu Rev Nutr 26, 1–22.

Carrico, C., Meyer, J.G., He, W., Gibson, B.W., and Verdin, E. (2018). The Mitochondrial Acylome Emerges: Proteomics, Regulation by Sirtuins, and Metabolic and Disease Implications. Cell Metab 27, 497–512.

Chayka, O., D’Acunto, C.W., Middleton, O., Arab, M., and Sala, A. (2015). Identification and pharmacological inactivation of the MYCN gene network as a therapeutic strategy for neuroblastic tumor cells. J Biol Chem 290, 2198–2212.

Choudhary, C., Kumar, C., Gnad, F., Nielsen, M.L., Rehman, M., Walther, T.C., Olsen, J.V., and Mann, M. (2009). Lysine acetylation targets protein complexes and co-regulates major cellular functions. Science 325, 834–840.

Colaert, N., Helsens, K., Martens, L., Vandekerckhove, J., and Gevaert, K. (2009). Improved visualization of protein consensus sequences by iceLogo. Nat Methods 6, 786–787.

Cox, J., and Mann, M. (2008). MaxQuant enables high peptide identification rates, individualized p.p.b.-range mass accuracies and proteome-wide protein quantification. Nat Biotechnol 26, 1367–1372.

Dickinson, M.E., Flenniken, A.M., Ji, X., Teboul, L., Wong, M.D., White, J.K., Meehan, T.F., Weninger, W.J., Westerberg, H., Adissu, H., et al. (2016). High-throughput discovery of novel developmental phenotypes. Nature 537, 508–514.

Ducker, G.S., and Rabinowitz, J.D. (2017). One-Carbon Metabolism in Health and Disease. Cell Metab 25, 27–42.

Furman, B.L. (2015). Streptozotocin-Induced Diabetic Models in Mice and Rats. Curr Protoc Pharmacol 70, 5 47 41–45 47 20.

Girgis, S., Nasrallah, I.M., Suh, J.R., Oppenheim, E., Zanetti, K.A., Mastri, M.G., and Stover, P.J. (1998). Molecular cloning, characterization and alternative splicing of the human cytoplasmic serine hydroxymethyltransferase gene. Gene 210, 315–324.

Grbesa, I., Kalo, A., Beluzic, R., Kovacevic, L., Lepur, A., Rokic, F., Hochberg, H., Kanter, I., Simunovic, V., Munoz-Torres, P.M., et al. (2017). Mutations in S-adenosylhomocysteine hydrolase (AHCY) affect its nucleocytoplasmic distribution and capability to interact with S-adenosylhomocysteine hydrolase-like 1 protein. Eur J Cell Biol 96, 579–590.

Hashim, S.A., and VanItallie, T.B. (2014). Ketone body therapy: from the ketogenic diet to the oral administration of ketone ester. J Lipid Res 55, 1818–1826.

Haws, S.A., Yu, D., Ye, C., Wille, C.K., Nguyen, L.C., Krautkramer, K.A., Tomasiewicz, J.L., Yang, S.E., Miller, B.R., Liu, W.H., et al. (2020). Methyl-Metabolite Depletion Elicits Adaptive Responses to Support Heterochromatin Stability and Epigenetic Persistence. Mol Cell 78, 210–223 e218.

Huang da, W., Sherman, B.T., and Lempicki, R.A. (2009). Systematic and integrative analysis of large gene lists using DAVID bioinformatics resources. Nat Protoc 4, 44–57.

Huang, H., Tang, S., Ji, M., Tang, Z., Shimada, M., Liu, X., Qi, S., Locasale, J.W., Roeder, R.G., Zhao, Y., et al. (2018a). p300-Mediated Lysine 2-Hydroxyisobutyrylation Regulates Glycolysis. Mol Cell 70, 663–678 e666.

Huang, H., Wang, D.L., and Zhao, Y. (2018b). Quantitative Crotonylome Analysis Expands the Roles of p300 in the Regulation of Lysine Crotonylation Pathway. Proteomics 18, e1700230.

Kaczmarska, Z., Ortega, E., Goudarzi, A., Huang, H., Kim, S., Marquez, J.A., Zhao, Y., Khochbin, S., and Panne, D. (2017). Structure of p300 in complex with acyl-CoA variants. Nat Chem Biol 13, 21–29.

Kang, H.B., Fan, J., Lin, R., Elf, S., Ji, Q., Zhao, L., Jin, L., Seo, J.H., Shan, C., Arbiser, J.L., et al. (2015). Metabolic Rewiring by Oncogenic BRAF V600E Links Ketogenesis Pathway to BRAF-MEK1 Signaling. Mol Cell 59, 345–358.

Katada, S., Imhof, A., and Sassone-Corsi, P. (2012). Connecting threads: epigenetics and metabolism. Cell 148, 24–28.

Koeslag, J.H., Noakes, T.D., and Sloan, A.W. (1980). Post-exercise ketosis. J Physiol 301, 79–90.

Lin, H., Su, X., and He, B. (2012). Protein lysine acylation and cysteine succination by intermediates of energy metabolism. ACS Chem Biol 7, 947–960.

Liu, K., Li, F., Sun, Q., Lin, N., Han, H., You, K., Tian, F., Mao, Z., Li, T., Tong, T., et al. (2019). p53 beta-hydroxybutyrylation attenuates p53 activity. Cell Death Dis 10, 243.

Manszewski, T., Szpotkowski, K., and Jaskolski, M. (2017). Crystallographic and SAXS studies of S-adenosyl-l-homocysteine hydrolase from Bradyrhizobium elkanii. IUCrJ 4, 271–282.

Maurer, G.D., Brucker, D.P., Bahr, O., Harter, P.N., Hattingen, E., Walenta, S., Mueller-Klieser, W., Steinbach, J.P., and Rieger, J. (2011). Differential utilization of ketone bodies by neurons and glioma cell lines: a rationale for ketogenic diet as experimental glioma therapy. BMC Cancer 11, 315.

McGarry, J.D., and Foster, D.W. (1980). Regulation of hepatic fatty acid oxidation and ketone body production. Annu Rev Biochem 49, 395–420.

Mews, P., Donahue, G., Drake, A.M., Luczak, V., Abel, T., and Berger, S.L. (2017). Acetyl-CoA synthetase regulates histone acetylation and hippocampal memory. Nature 546, 381–386.

Newman, J.C., Covarrubias, A.J., Zhao, M., Yu, X., Gut, P., Ng, C.P., Huang, Y., Haldar, S., and Verdin, E. (2017). Ketogenic Diet Reduces Midlife Mortality and Improves Memory in Aging Mice. Cell Metab 26, 547–557 e548.

Newman, J.C., and Verdin, E. (2017). beta-Hydroxybutyrate: A Signaling Metabolite. Annu Rev Nutr 37, 51–76.

Nishida, Y., Rardin, M.J., Carrico, C., He, W., Sahu, A.K., Gut, P., Najjar, R., Fitch, M., Hellerstein, M., Gibson, B.W., et al. (2015). SIRT5 Regulates both Cytosolic and Mitochondrial Protein Malonylation with Glycolysis as a Major Target. Mol Cell 59, 321– 332.

Pajares, M.A., and Perez-Sala, D. (2006). Betaine homocysteine S-methyltransferase: just a regulator of homocysteine metabolism? Cell Mol Life Sci 63, 2792–2803.

Peleg, S., Feller, C., Ladurner, A.G., and Imhof, A. (2016). The Metabolic Impact on Histone Acetylation and Transcription in Ageing. Trends Biochem Sci 41, 700–711.

Pissios, P., Hong, S., Kennedy, A.R., Prasad, D., Liu, F.F., and Maratos-Flier, E. (2013). Methionine and choline regulate the metabolic phenotype of a ketogenic diet. Mol Metab 2, 306–313.

Poff, A.M., Ari, C., Arnold, P., Seyfried, T.N., and D’Agostino, D.P. (2014). Ketone supplementation decreases tumor cell viability and prolongs survival of mice with metastatic cancer. Int J Cancer 135, 1711–1720.

Puchalska, P., and Crawford, P.A. (2017). Multi-dimensional Roles of Ketone Bodies in Fuel Metabolism, Signaling, and Therapeutics. Cell Metab 25, 262–284.

Rardin, M.J., He, W., Nishida, Y., Newman, J.C., Carrico, C., Danielson, S.R., Guo, A., Gut, P., Sahu, A.K., Li, B., et al. (2013a). SIRT5 regulates the mitochondrial lysine succinylome and metabolic networks. Cell Metab 18, 920–933.

Rardin, M.J., Newman, J.C., Held, J.M., Cusack, M.P., Sorensen, D.J., Li, B., Schilling, B., Mooney, S.D., Kahn, C.R., Verdin, E., et al. (2013b). Label-free quantitative proteomics of the lysine acetylome in mitochondria identifies substrates of SIRT3 in metabolic pathways. Proc Natl Acad Sci U S A 110, 6601–6606.

Reed, W.D., and Ozand, P.T. (1980). Enzymes of L-(+)-3-hydroxybutyrate metabolism in the rat. Arch Biochem Biophys 205, 94–103.

Ringel, A.E., Tucker, S.A., and Haigis, M.C. (2018). Chemical and Physiological Features of Mitochondrial Acylation. Mol Cell 72, 610–624.

Roberts, M.N., Wallace, M.A., Tomilov, A.A., Zhou, Z., Marcotte, G.R., Tran, D., Perez, G., Gutierrez-Casado, E., Koike, S., Knotts, T.A., et al. (2017). A Ketogenic Diet Extends Longevity and Healthspan in Adult Mice. Cell Metab 26, 539–546 e535.

Robinson, A.M., and Williamson, D.H. (1980). Physiological roles of ketone bodies as substrates and signals in mammalian tissues. Physiol Rev 60, 143–187.

Ruan, H.B., and Crawford, P.A. (2018). Ketone bodies as epigenetic modifiers. Curr Opin Clin Nutr Metab Care 21, 260–266.

Sabari, B.R., Tang, Z., Huang, H., Yong-Gonzalez, V., Molina, H., Kong, H.E., Dai, L., Shimada, M., Cross, J.R., Zhao, Y., et al. (2015). Intracellular crotonyl-CoA stimulates transcription through p300-catalyzed histone crotonylation. Mol Cell 58, 203–215.

Sabari, B.R., Tang, Z., Huang, H., Yong-Gonzalez, V., Molina, H., Kong, H.E., Dai, L., Shimada, M., Cross, J.R., Zhao, Y., et al. (2018). Intracellular Crotonyl-CoA Stimulates Transcription through p300-Catalyzed Histone Crotonylation. Mol Cell 69, 533.

Sabari, B.R., Zhang, D., Allis, C.D., and Zhao, Y. (2017). Metabolic regulation of gene expression through histone acylations. Nat Rev Mol Cell Biol 18, 90–101.

Sahar, S., Masubuchi, S., Eckel-Mahan, K., Vollmer, S., Galla, L., Ceglia, N., Masri, S., Barth, T.K., Grimaldi, B., Oluyemi, O., et al. (2014). Circadian control of fatty acid elongation by SIRT1 protein-mediated deacetylation of acetyl-coenzyme A synthetase 1. J Biol Chem 289, 6091–6097.

Sato, S., Solanas, G., Peixoto, F.O., Bee, L., Symeonidi, A., Schmidt, M.S., Brenner, C., Masri, S., Benitah, S.A., and Sassone-Corsi, P. (2017). Circadian Reprogramming in the Liver Identifies Metabolic Pathways of Aging. Cell 170, 664–677 e611.

Serefidou, M., Venkatasubramani, A.V., and Imhof, A. (2019). The Impact of One Carbon Metabolism on Histone Methylation. Front Genet 10, 764.

Smith, A.C., and Robinson, A.J. (2016). MitoMiner v3.1, an update on the mitochondrial proteomics database. Nucleic Acids Res 44, D1258–1261.

Stender, S., Chakrabarti, R.S., Xing, C., Gotway, G., Cohen, J.C., and Hobbs, H.H. (2015). Adult-onset liver disease and hepatocellular carcinoma in S-adenosylhomocysteine hydrolase deficiency. Mol Genet Metab 116, 269–274.

Tan, M., Peng, C., Anderson, K.A., Chhoy, P., Xie, Z., Dai, L., Park, J., Chen, Y., Huang, H., Zhang, Y., et al. (2014). Lysine glutarylation is a protein posttranslational modification regulated by SIRT5. Cell Metab 19, 605–617.

Tognini, P., Murakami, M., Liu, Y., Eckel-Mahan, K.L., Newman, J.C., Verdin, E., Baldi, P., and Sassone-Corsi, P. (2017). Distinct Circadian Signatures in Liver and Gut Clocks Revealed by Ketogenic Diet. Cell Metab 26, 523–538 e525.

Veech, R.L. (2014). Ketone ester effects on metabolism and transcription. J Lipid Res 55, 2004–2006.

Verdin, E., and Ott, M. (2015). 50 years of protein acetylation: from gene regulation to epigenetics, metabolism and beyond. Nat Rev Mol Cell Biol 16, 258–264.

Vizcaino, J.A., Csordas, A., Del-Toro, N., Dianes, J.A., Griss, J., Lavidas, I., Mayer, G., Perez-Riverol, Y., Reisinger, F., Ternent, T., et al. (2016). 2016 update of the PRIDE database and its related tools. Nucleic Acids Res 44, 11033.

Wagner, G.R., Bhatt, D.P., O’Connell, T.M., Thompson, J.W., Dubois, L.G., Backos, D.S., Yang, H., Mitchell, G.A., Ilkayeva, O.R., Stevens, R.D., et al. (2017). A Class of Reactive Acyl-CoA Species Reveals the Non-enzymatic Origins of Protein Acylation. Cell Metab 25, 823–837 e828.

Wagner, G.R., and Hirschey, M.D. (2014). Nonenzymatic protein acylation as a carbon stress regulated by sirtuin deacylases. Mol Cell 54, 5–16.

Walker, A.K. (2017). 1-Carbon Cycle Metabolites Methylate Their Way to Fatty Liver. Trends Endocrinol Metab 28, 63–72.

Wang, Y., Kavran, J.M., Chen, Z., Karukurichi, K.R., Leahy, D.J., and Cole, P.A. (2014). Regulation of S-adenosylhomocysteine hydrolase by lysine acetylation. J Biol Chem 289, 31361–31372.

Weber, D.D., Aminazdeh-Gohari, S., and Kofler, B. (2018). Ketogenic diet in cancer therapy. Aging (Albany NY) 10, 164–165.

Xie, Z., Zhang, D., Chung, D., Tang, Z., Huang, H., Dai, L., Qi, S., Li, J., Colak, G., Chen, Y., et al. (2016). Metabolic Regulation of Gene Expression by Histone Lysine beta-Hydroxybutyrylation. Mol Cell 62, 194–206.

Yeo, E.J., and Wagner, C. (1992). Purification and properties of pancreatic glycine N-methyltransferase. J Biol Chem 267, 24669–24674.

Zhao, S., Xu, W., Jiang, W., Yu, W., Lin, Y., Zhang, T., Yao, J., Zhou, L., Zeng, Y., Li, H., et al. (2010). Regulation of cellular metabolism by protein lysine acetylation. Science 327, 1000–1004.

